# Microbial tryptophan catabolism affects the vector competence of *Anopheles*

**DOI:** 10.1101/2021.02.15.431262

**Authors:** Yuebiao Feng, Yeqing Peng, Han Wen, Xiumei Song, Yanpeng An, Huiru Tang, Jingwen Wang

## Abstract

The influence of microbiota on mosquito physiology and vector competence is becoming increasingly clear but our understanding of interactions between microbiota and mosquitoes still remains incomplete. Here we show that gut microbiota of *Anopheles stephensi*, a competent malaria vector, participates mosquito tryptophan metabolism. Elimination of microbiota by antibiotics treatment leads to the accumulation of tryptophan (Trp) and its metabolites, kynurenine (Kyn), 3‐hydroxykynurenine (3‐HK) and xanthurenic acid (XA). Of these, 3‐HK impairs the structure of peritrophic matrix (PM), thereby promoting *Plasmodium berghei* infection. Among the major gut microbiota in *An. stephensi, Pseudomonas alcaligenes* plays a role in catabolizing 3‐HK as revealed by whole genome sequencing and LC‐MS metabolic analysis. The genome of *P. alcaligenes* encodes kynureninase (KynU) that is responsible for the conversion of 3‐HK to 3‐Hydroxyanthranilic acid (3‐HAA). Mutation of this gene abrogates the ability of *P. alcaligenes* to metabolize 3‐HK, which in turn abolishes its role on PM protection. Colonization of *An. stephensi* with KynU mutated *P. alcaligenes* fails to protect mosquitoes against parasite infection as effectively as those with wild type bacterium. In summary, we identify an unexpected function of gut microbiota in controlling mosquito tryptophan metabolism with the major consequences on vector competence.

## Introduction

*Anopheles* mosquitoes, the primary vectors of malaria, are colonized by a population of diverse and dynamic microbiota^1‐3^. These microbes play important roles on several key physiological mosquito functions, including development, nutrition and vector competence^4^. Of these, microbiota is vital in promoting blood meal digestion that provides essential nutrients for egg development^5^. Elimination of these microbes impairs blood cell lysis and slows down protein catabolism^5^. So far only a limited number of commensal bacteria involved in blood proteolysis have been characterized. *Serratia marcescens* contributes to erythrocytes lysis by producing hemolysins in *Anopheles* mosquito^6^. *Elizabethkingia anopheles* possesses the heme-binding protein, HemS, that oxidatively cleaves heme to biliverdin^7^. *Acinetobacter* isolates in *Aedes albopictus* are able to metabolize blood component, α‐keto‐valeric acid and glycine, and improve blood digestion^8^. Proteins, accounting for about 95% of the blood constituents, are the primary source of amino acids for mosquitoes^9^. There is still little mechanistic insight into the contribution of microbiota toward mosquito amino acid metabolism.

Tryptophan (Trp) is one of the essential amino acids that mainly supplied by ingested blood meal^10^. In addition to be used in protein synthesis, tryptophan is oxidized through kynurenine pathway, resulting in the production of kynurenine (Kyn), 3‐hydroxykynurenine (3‐HK) and xanthurenic acid (XA), and transformed into 5‐Hydroxy‐L‐tryptophan (5‐HT, serotonin) and derivatives through serotonin pathway^11,12^. The tryptophan metabolites play vital roles in various physiological processes of mosquitoes. 3‐HK is the precursor for production of eye pigmentation during pupal development^12^. Serotonin that functions as a neurohormone in mosquitoes modulates insulin‐like peptides expression^13^, ion transportation^14^, feeding behavior^15^, salivation^16^ and heart rate^17^. In mammalians, tryptophan and its metabolites are also key mediators that regulate immune responses and gut barrier function, thereby affecting hosts’ susceptibility to pathogen infections^18‐23^. However, little is known about the influence of mosquito tryptophan metabolism on pathogen transmission.

In this study, we show that microbiota in *Anopheles stephensi*, participates mosquito tryptophan metabolism. Elimination of microbiota leads to the accumulation of multiple tryptophan (Trp) catabolites belong to kynurenine pathway. Among these metabolites, 3‐hydroxykynurenine (3‐HK) elevation impairs the integrity of peritrophic matrix that functions as a physical barrier in the midgut, and facilitates *Plasmodium berghei* infection. The gut commensal bacterium, *P. alcaligenes*, processes the enzyme Kynureninase (KynU) that is responsible for catabolizing 3‐HK. Mutation of KynU abolished the capacity of *P. alcaligenes* to degrade 3‐HK and reduces its inhibitory effect on *P. berghei*. Collectively, our results demonstrate that a direct cross‐talk of tryptophan metabolism between *An. stephensi* and gut microbiota controls the outcome of parasite infections.

## Result

### Microbiota participates Trp metabolism

To determine whether microbiota participates mosquito Trp metabolism, we performed a targeted metabolomics analysis by liquid chromatography–mass spectrometry (LC‐MS). Tryptophan and its metabolites were analyzed in normal and antibiotics treated (Abx) *An. stephensi* prior to (0 h) and 24 h post blood meal. Totally 15 Trp metabolites were detected (Fig. 1a,Supplementary Table 1). Trp and the metabolites of kynurenine pathway (KP), Kyn, 3‐HK and XA, were significantly more abundant in Abx mosquitoes than those in controls prior to blood feeding (Fig. 1b). 5‐HT and the bacterial derived metabolites 3‐HAA were reduced, while indole‐3‐aldehyde (IAld) were increased significantly in Abx mosquitoes 24 h post blood meal (Extended Data Fig. 1). These results indicate that microbiota modulates tryptophan metabolism in *An. stephensi*.

**Fig. 1.**
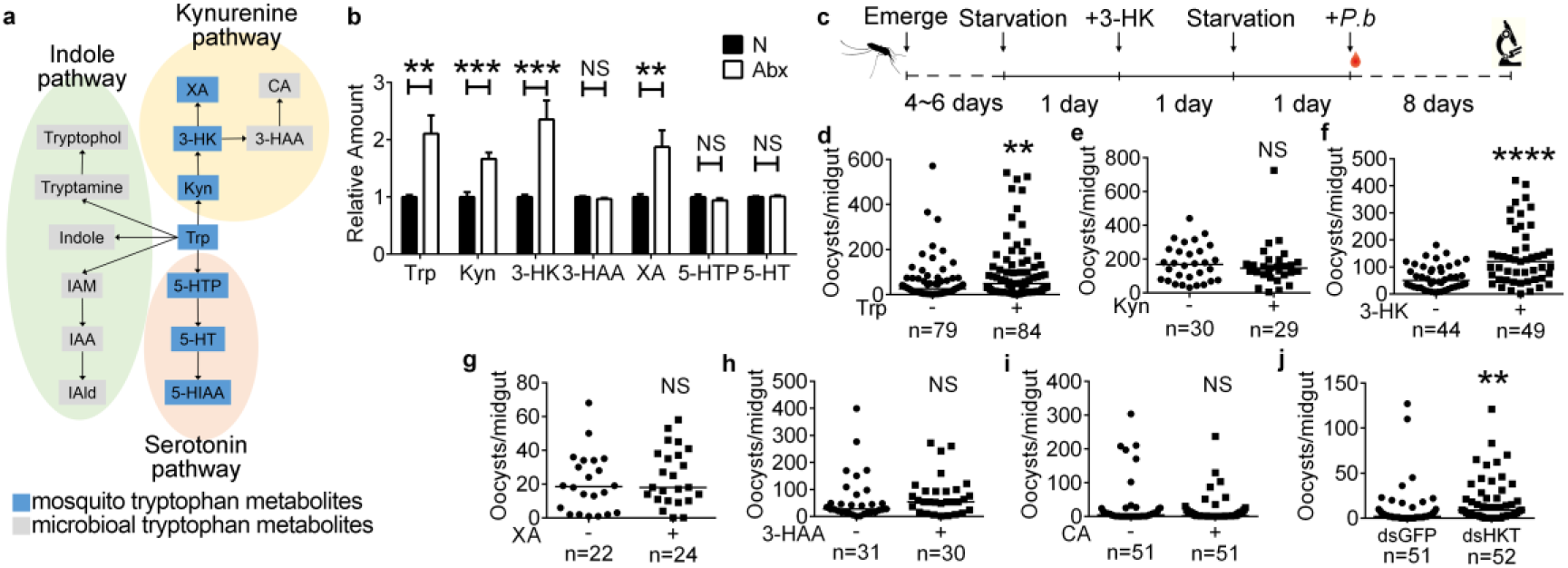
Microbiota regulated Trp metabolism modulates *P. berghei* infection. **a,** Standards associated with tryptophan metabolism in this study. Trp, Tryptophan; Kyn, Kynurenine; 3‐HK, 3‐Hydroxy‐kynurenine; XA, Xanthurenic acid; 3‐HAA, 3‐Hydroxyanthranilic acid; CA, Cinnabarinic acid; 5‐HTP, 5‐ Hydroxy‐tryptophan; 5‐HT, 5‐Hydroxytryptamine; 5‐HIAA, 5‐ Hydroxyindoleacetate, IAM, Indole‐3‐acetamide; IAA, Indole‐3‐acetic acid; IAld, Indole‐3‐acetaldehyde. **b**, The relative amount of Trp, Kyn, 3‐HK, XA, 3‐HAA, 5‐HT and 5‐HTP in normal (N, n=10) and antibiotics‐treated (Abx, n=9) mosquitoes prior to blood meal. Error bars indicate standard errors. **c**, Workflow of Trp metabolite treatments on *An. stephensi*. **d‐i**, Influence of Trp metabolites, Trp (**d**), Kyn (**e**), 3‐HK (**f**), XA (**g**), 3‐HAA (**h**) and CA (**i**) on *P. berghei* infection. **j**, Influence of *HKT* knockdown on *Plasmodium* infection. Data were pooled from two independent experiments. Horizontal black bars indicate the median values. Significance was determined by Student’s t‐test in (**b**) and by Mann‐ Whitney tests in (**d‐j**). **P<0.01, ***P<0.001, ****P<0.0001.

Mosquitoes of which microbiota is removed are more susceptible to *Plasmodium* infection^24^. In light of our finding that the levels of KP associated compounds, Trp, Kyn, 3‐HK, and XA, and 3‐HAA were affected by microbiota, we hypothesized that these five KP metabolites might contribute to the increased susceptibility of mosquitoes to *P. berghei* infection. We next orally administrated these five metabolites to *An. stephensi* for 24 h, followed by fed on mice infected with *P. berghei* 1 day later. Oocyst number was examined 8‐ day post infection (Fig. 1c). The amount of tryptophan was used as described^25^ and the amounts of Kyn, 3‐HK, XA, and 3‐HAA were used based on their corresponding level in normal mosquitoes as revealed by LC‐MS analysis (Supplementary Table 2 and Table 3). As expected, orally supplementation of Trp and 3‐HK via sugar meal both significantly increased oocysts number in *An. stephensi* (Fig. 1d, f). However, administration of Kyn, XA, and 3‐HAA, had no influence on *P. berghei* infection (Fig. 1e, g, h). In addition, we examined the influence of cinnabarinic acid (CA), the downstream product of 3‐HAA, on *P. berghei* infection. No difference of oocyst number was observed between CA supplemented and non‐supplemented groups (Fig. 1i). To further confirm that 3‐HK affects *P. berghei* infection, we knocked down the gene encoding mosquito 3‐hydroxykynurenine transaminase (HKT), which catalyzes the conversion of 3‐HK into XA, and analyzed mosquito infection rate. Knocking down *HKT* (dsHKT) significantly increases oocyst number compared to dsGFP controls (Fig. 1j). Altogether, these results suggest that tryptophan metabolism, especially kynurenine pathway is under the regulation by both mosquito and microbiota. The accumulation of 3‐HK contributes to the increased susceptibility to parasite infection.

### 3‐HK accumulation impairs PM integrity

Due to the high level of 3‐HK in mosquitoes (Supplementary Table 2) and its capacity to generate oxidative stress^26,27^, we hypothesized that the elevation of 3‐HK might play a major role in disturbing the redox homeostasis in the midgut, thereby impairing the midgut epithelial barrier function and facilitating parasite infection. We first examined the ROS level in the midgut of *An. stephensi* supplemented with or without 3‐HK at 0 and 24 h post blood feeding by dihydroethidium (DHE) staining. In contrast to our hypothesis, no difference of ROS level was observed between 3‐HK treated and control mosquitoes either prior to or post blood meal (Extended Data Fig. 2). Midgut epithelial cells turn over rapidly due to damage from digestion and toxins^28^. We next examined whether ingestion of 3‐HK could damage midgut epithelial cell via staining the apoptotic cells and the mitotic intestinal stem cells^28^. Again, 3‐HK treatment didn’t elicit significant changes in midgut epithelial cells, compared with controls (Extended Data Fig. 3, 4).

**Fig. 2.**
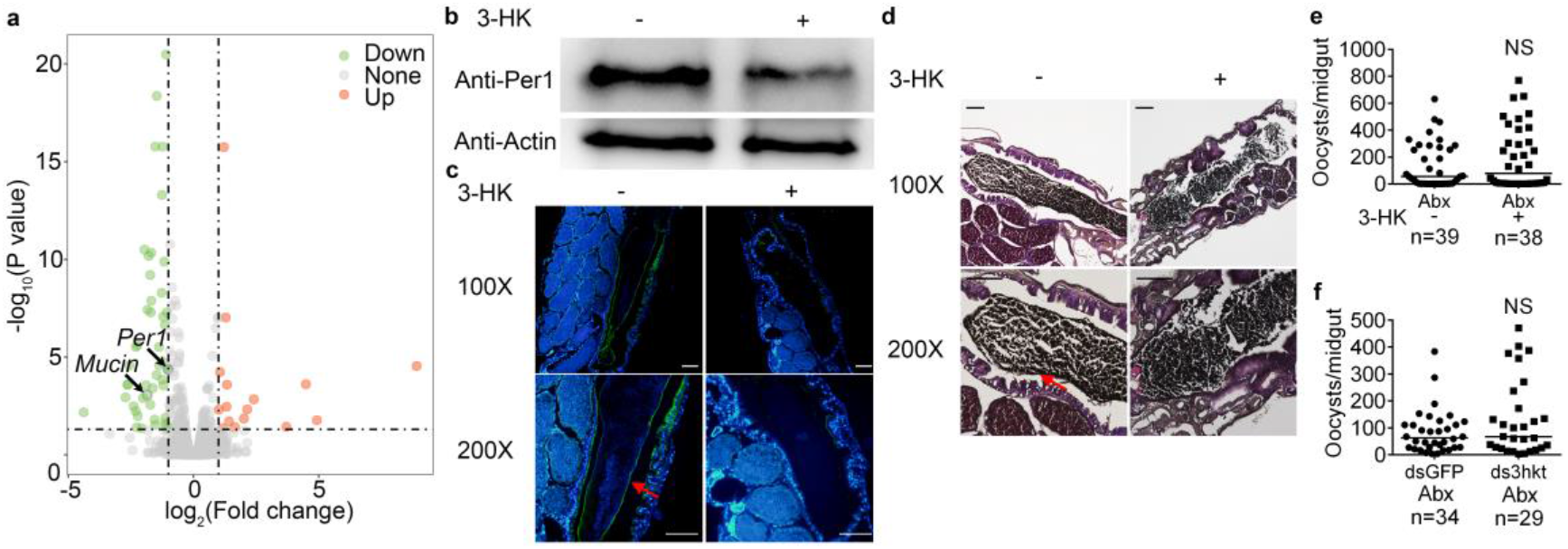
3‐HK promotes *Plasmodium* infection via impairing PM. **a,** Volcano plot shows differentially expressed genes in the midguts of mosquitoes fed with/without 3‐HK at 24 h post infection (hpi). Upregulated genes are shown in orange; downregulated genes are shown in green. **b**, Western blot of Per1 in the midgut of 3‐HK‐treated (+) mosquitoes and control (‐) 24 h post normal blood meal. **c**, Immunostaining of Per1 (green) in the midgut of 3‐HK‐treated (+) and control (‐) mosquitoes at 100× and 200× magnification. Nuclei were stained with DAPI (blue). Red arrows indicate Per1. Images are representative of at least five midguts. Scale bars represent 100 µm. **d**, PAS staining of PM structure in 3‐HK‐treated (+) and control (‐) mosquitoes at 100× and 200× magnification. Red arrows indicate the PM structure. Images are representative of at least four individual mosquito midguts. Scale bars represent 100 µm. **e**, Oocyst numbers of Abx (‐) and Abx mosquitoes supplemented with 3‐HK (+). **f**, Oocyst numbers of Abx mosquitoes treated with dsHKT and dsGFP. Data were pooled from two independent experiments (**e, f**). Each dot represents an individual mosquito. Horizontal black bars indicate the median values. Significance was determined by Mann‐Whitney tests.

**Fig. 3.**
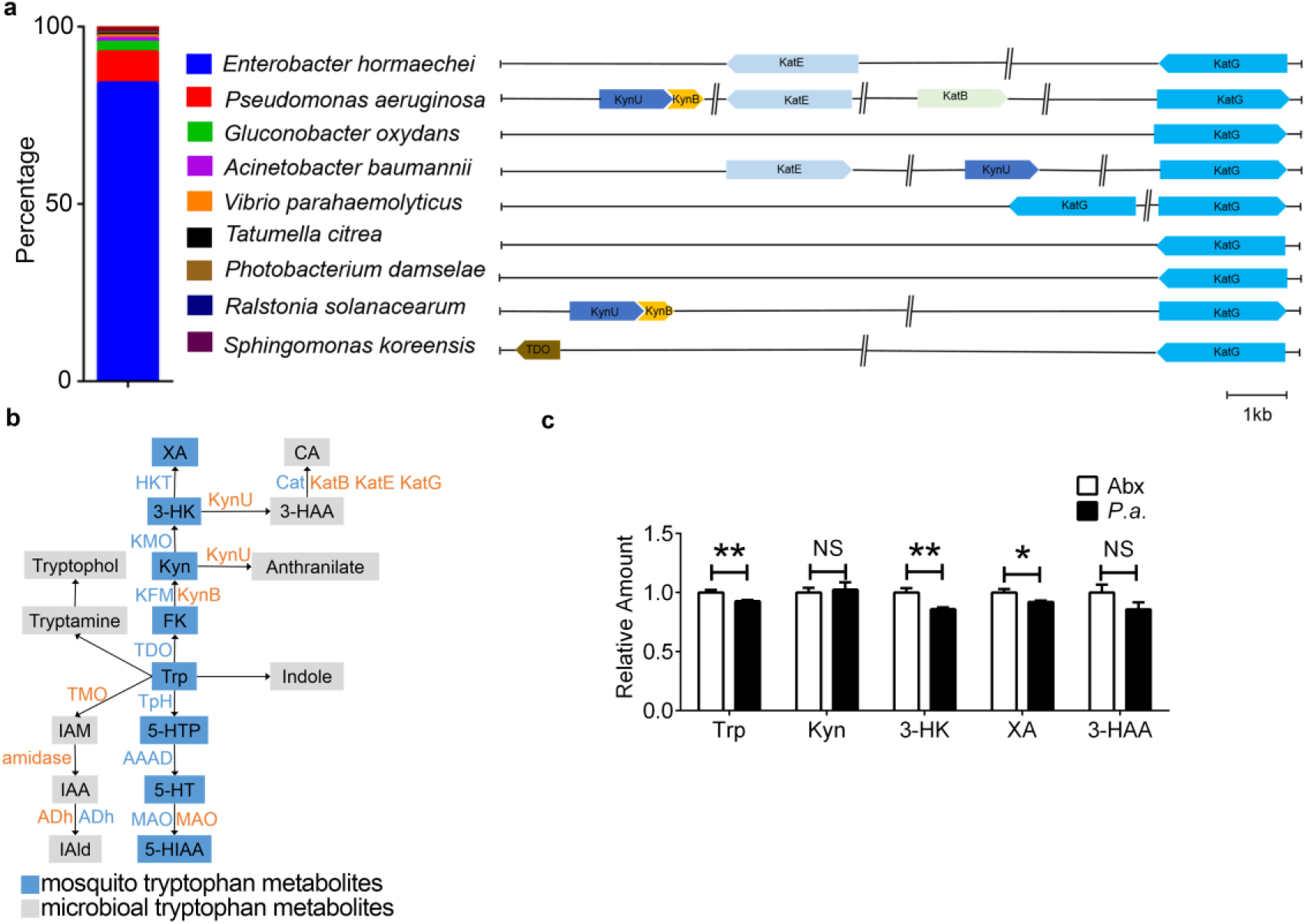
Commensal *P. alcaligenes* catabolizes mosquito 3‐HK. **a,** Kynurenine pathways of major commensal bacteria in *An. stephensi*. Left panel, relative abundance of major bacteria genera in lab‐reared mosquito by 16S rRNA pyrosequencing. The column represents six pooled midguts. Right panel, gene clusters of kynurenine metabolic pathways in the representative bacteria species of each genus. **b**, Overview of Trp catabolism through mosquito (blue) and *P. alcaligenes* (orange) pathways. HKT, 3‐ hydroxykynurenine; KMO, Kynurenine 3‐Monooxygenase; KFM, kynurenine formamidase; TDO, Tryptophan 2,3‐Dioxygenase; TpH, Tryptophan Hydroxylase; AAAD, Aromatic Amino Acid Decarboxylase; MAO, Monoamine Oxydase; KynU, kynureninase; Cat, Catalase; Kat, Peroxidases; TMO, Tryptophan 2‐monooxygenase; ADh, aldehyde dehydrogenase. **c**, The relative amounts of Trp metabolites in *P. alcaligenes* recolonized (*P*.*a*.) and antibiotics treated mosquito (Abx) before blood meal. Error bars indicate standard errors (n=9). Significance was determined by Student’s *t*‐test. * P<0.05, **P<0.01.

**Fig. 4.**
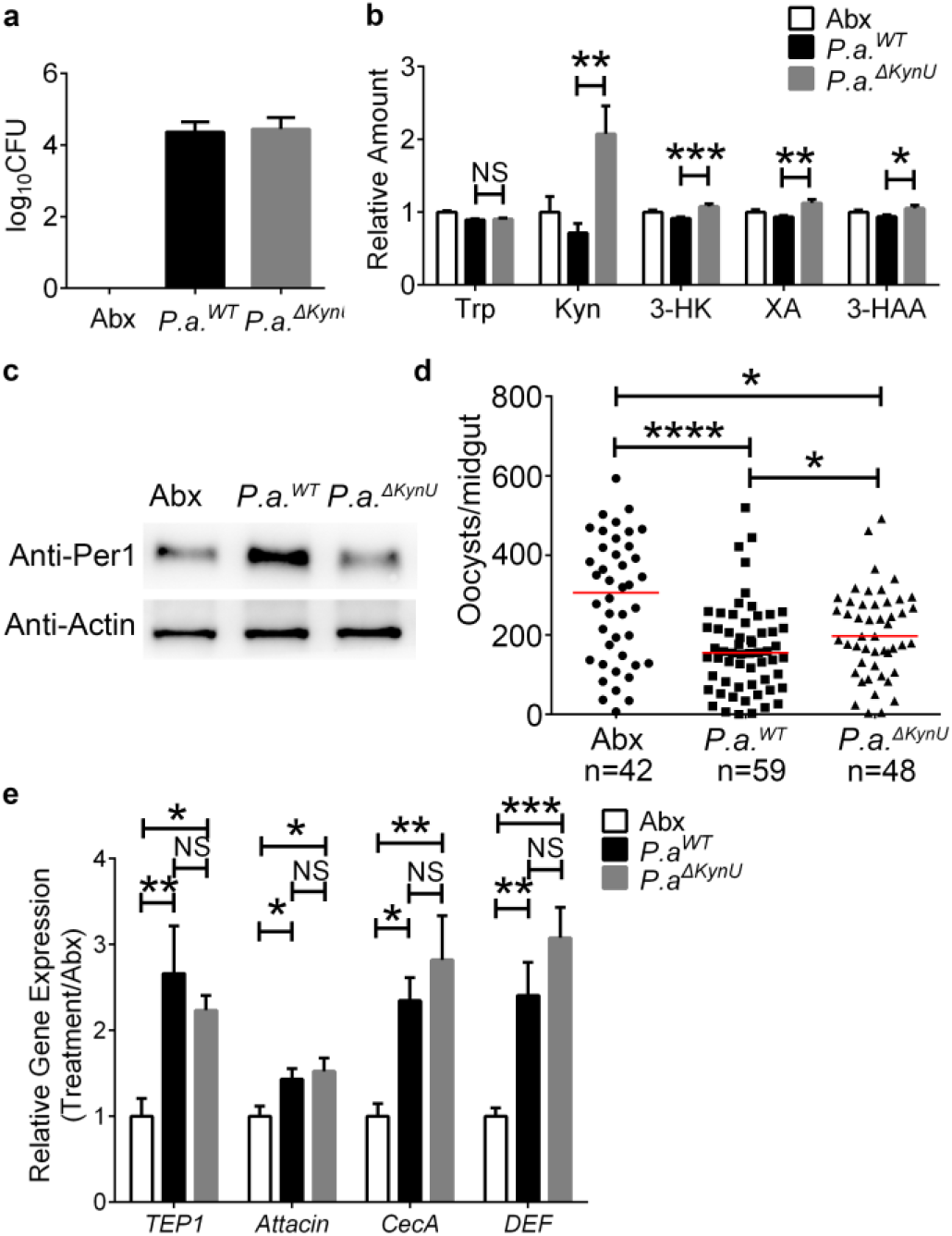
*P*.*alcaligenes* KynU degrades 3‐HK in mosquitoes. **a,** Midgut bacterial loads in Abx and Abx mosquitoes colonized with *P*.*a*.^*WT*^ and *P*.*a*.*Δ*^*KynU*^ before blood meal (n=6). **b**, The relative amounts of Trp metabolites in Abx (n=10), and Abx mosquitoes colonized with *P*.*a*.^*WT*^ and *P*.*a*.*^ΔKynU^* (n=12) before blood meal. Error bars indicate standard errors. **c**, Western blot of Per1 in the midgut of Abx, *P*.*a*.^*WT*^ and *P*.*a*.^*ΔKynU*^ recolonized mosquitoes 24 h post normal blood meal. Images are representative of three independent experiments. **d**, Oocyst numbers of Abx, *P*.*a*.^*WT*^ and *P*.*a*.^*ΔKynU*^ recolonized mosquitoes. Data were pooled from two independent experiments. Horizontal black bars indicate the median values. e, Relative expression levels of immune related genes in Abx, *P*.*a*.^*WT*^ and *P*.*a*.^*ΔKynU*^ recolonized mosquitoes. The expression level of target gene was normalized to *S7*. The relative expression level of immune genes in *P*.*a*.^*WT*^ and *P*.*a*.^*ΔKynU*^colonized mosquitoes was normalized to the gene’s expression in Abx, respectively. Error bars indicate standard errors (n = 6∼8). Results from one of two independent experiments are shown. Significance was determined by ANOVA tests in (**b, d, e**). * P<0.05, **P<0.01, ***P<0.001, ****P<0.0001.

To further investigate the mechanism that 3‐HK facilitates *P. berghei* infection, RNA sequencing was performed on midguts of mosquitoes treated with or without 3‐HK at 24 h post an infectious blood meal. There were only 172 genes differentially expressed with 47 genes upregulated and 125 downregulated in 3‐HK treated mosquitoes (Supplementary Table 4). Among the downregulated genes with fold changes >2, we identified two genes associated with midgut epithelial barrier function, including *mucin* and *peritrophin1* (Fig. 2a). Peritrophin and mucin are important components of peritrophic matrix (PM) that protects midgut from pathogens invasion, abrasion and toxic compounds^29‐33^. To examine whether orally administration of 3‐HK could impair PM, we analyzed Peritrophin 1 (Per1) protein level and PM structure in 3‐HK supplemented mosquitoes. In agreement with the RNA‐seq results, supplementation of 3‐HK dramatically reduced Per1 protein level in the midguts (Fig 2b, c), and impaired the PM structure (Fig. 2d).

To further confirm that 3‐HK‐mediated increased parasite infection relies on the integrity of PM, we disrupted the formation of PM by elimination of gut microbiota as described^32,33^, and examined the influence of 3‐HK on *Plasmodium* infection. Orally administration of 3‐HK to PM compromised mosquitoes (Abx) failed to increase the number of oocysts compared with non‐ 3‐HK treated ones (Fig. 2e). Consistently, knockdown of HKT has no influence on infection outcome when PM was impaired (Fig. 2f). We next performed the same analyses on mosquitoes supplemented with Trp. Again, Trp oral administration reduced protein level of Per1 compared to controls (Extended Data Fig. 5a). Trp supplementation no longer affected *P. berghei* infection when PM was absent (Extended Data Fig. 5b). Altogether, our results suggest that elevation of 3‐HK impairs PM integrity, which in turn promotes *P. berghei* infection.

### *Pseudomonas alcaligenes* catabolizes 3‐HK

Given that gut microbiota prevents the accumulation of PM impairing 3‐HK, we were interested to identify which gut commensal bacterium plays a major role in 3‐HK catabolism. The population structure of gut microbiota was analyzed in the laboratory reared *An. stephensi* before blood meal. The top nine abundant genera were *Enterobacter, Pseudomonas, Gluconobacter, Acinetobacter, Vibrio, Tatumella, Photobacterium, Ralstonia* and *Sphingomaoans* (Fig. 3a). We next analyzed the genome sequence of the representative species in Kyoto Encyclopedia of Genes and Genomes database to evaluate their genetic capacity to catabolize 3‐HK. Bacteria from Genera *Pseudomonas* and *Ralstonia* both contain kynurenine formamidase (KynB) that is responsible for the production of Kyn from N‐formylkynurenine (FK), and the kynureninase (KynU) that hydrolyzes Kyn and 3‐HK to anthranilate and 3‐HAA, respectively (Fig. 3a)^34,35^. As *Pseudomonas* is the second abundant genus in *An. stephensi*, we next investigated the role of *Pseudomonas* in 3‐HK metabolism. The major *Pseudomonas* species in our colony *An. stephensi* is *Pseudomonas alcaligenes* (Extended Data Fig. 6a).

To examine the genetic capacity of the commensal *P. alcaligenes* on Trp catabolism, we obtained complete genome sequence of this bacterium by Illumina sequencing. We identified seven genes that were associated with Trp catabolism in *P. alcaligenes*. The three enzymes, kynurenine formamidase (encoded by *kynB*) and kynureninase (encodey by *kynU*) and one peroxidase (encoded by *kat*) were associated with Trp catabolism through kynurenine pathway (Fig. 3b). Tryptophan 2‐monooxygenase (encoded by *TMO*), amidase, and aldehyde dehydrogenase (encoded by *ADh*) were associated with Indole pathway, and monoamine oxidase (encoded by *MAO*) was responsible for the conversion from 5‐HT to 5‐HIAA (Fig. 3b).

To validate the ability of *P. alcaligenes* to catabolize 3‐HK in mosquitoes, we compared the Trp metabolism between Abx and Abx mosquitoes re‐colonized with *P. alcaligenes* by LC‐MS. *P. alcaligenes* reached 1.2×10^4^ CFU/midgut two days post inoculation (Extended Data Fig. 6b). As expected, re‐colonization of *P. alcaligenes* in Abx mosquitoes significantly reduced the levels of Trp, 3‐HK, and XA, compared with those in Abx mosquitoes (Fig 3c). However, we failed to detect cinnabarinic acid (CA), the end product of 3‐HK, *in vivo* possibly due to their low level. Altogether, these results confirmed the commensal bacterium *P. alcaligenes* is responsible for metabolizing 3‐HK in *An. stephensi*.

### Bacterial Kynureninase catabolizes 3‐HK

Kynureninase (KynU) is annotated to be responsible for the catabolism of 3‐HK in *P. alcaligenes* (Fig. 3b). To verify the role of KynU on 3‐HK degradation, we generated a KynU mutant, *P*.*a*.^*ΔKynU*^, lacking a 1005‐bp coding sequence of KynU (Extended Data Fig. 7a). Mutation of KynU didn’t influence bacterial growth either *in vivo* or *in vitro* (Fig. 4a, Extended Data Fig. 7b). We again assessed the influence of KynU on tryptophan metabolism in *An. stephensi*. Metabolites associated with KP were examined in Abx mosquitoes, and Abx mosquitoes colonized with wild type *P. alcaligenes, P*.*a*.^*WT*^ and KynU mutant, *P*.*a*. ^*Δ KynU*^. Mutation of KynU abolished the capability of *P. alcaligenes* to catabolize 3‐HK as mosquitoes re‐colonized with *P*.*a*.^*ΔKynU*^ had significantly higher level of 3‐HK than those re‐colonized with *P*.*a*.^*WT*^ (Fig. 4b). Furthermore, the levels of Kyn, XA and 3‐HAA, were accumulated in *P*.*a*.^*ΔKynU*^ colonized mosquitoes compared to *P*.*a*.*WT* colonized ones (Fig. 4b). Mutation of KynU didn’t influence the overall Trp metabolic activity of mosquitoes because the Trp abundance is comparable between *P*.*a*.^*WT*^ and *Pa*^*ΔKynU*^ colonized ones (Fig. 4b). As Trp is co‐metabolized by mosquito and its microbiota, blocking KynU activity in *P. alcaligenes* might increase the Trp metabolic activities of mosquito through indole and 5‐HT pathways.

We next examined the influence of bacterial KynU mutation on mosquito PM formation using Per1 as an indicator by western blot. As expected, re‐ colonization of *P*.*a*.^*ΔKynU*^ failed to induce Per1 protein expression as *P*.*a*.^*WT*^ did (Fig 4c). The inhibitory effect of *P. alcaligenes* on *P. berghei* was decreased significantly when KynU was mutated (Fig. 4d). However, colonization of these mutant bacteria still increased resistance of mosquitoes to parasite infection compared to microbiota cleared ones (Abx). It might due to the stimulation of *Pa*^*ΔKynU*^ in mosquito immune system. We next compared the expression level of four immune related genes, including *TEP1, CecA, GAM* and *DEF*, between *P*.*a*.^*WT*^ and *P*.*a*.^*ΔKynU*^ re‐colonized mosquitoes 24 hpi. The expression levels of all four immune genes were comparable between *P*.*a*.^*WT*^ and *P*.*a*.^*ΔKynU*^ colonized mosquitoes (Fig. 4e). Altogether, these results suggest that in addition to stimulating mosquito immune responses, *P. alcaligenes* inhibits *Plasmodium* infection through participating Trp catabolism to prevent the accumulation of PM damaging 3‐HK

## Discussion

Mosquito microbiota is known to inhibit malaria parasite infection through strengthening the immune system, stimulating the synthesis of peritrophic matrix (PM), and secreting anti‐plasmodial metabolites^1,3,36,37^. We have established a previously unknown role for gut microbiota in promoting antiparasitic responses through participating mosquito tryptophan metabolism. Elimination of microbiota by antibiotics treatment leads to the accumulation multiple metabolites of tryptophan. Among these metabolites, 3‐HK plays a role in impairing PM structure, thereby increasing susceptibility of mosquitoes to *P. berghei*. A midgut commensal bacterium, *P. alcaligenes*, is responsible for protecting mosquitoes against *P*.*berghei* infection by catabolizing 3‐HK. Mutation of KynU, the 3‐HK catabolizing enzyme abolishes the capability of *P. alcaligenes* to degrade 3‐HK, which in turn facilitates parasite infection.

The study of metabolic interactions between insects and microbiota have been focused on the role of their obligate symbionts in provision of dietary limited nutrients. For example, obligate haematophagous arthropods rely on their endosymbionts to provide B vitamins and cofactors that are scarce in animal blood^38^. The herbivorous insects obtain essential amino acids that are deficient in the plant sap from their endosymbionts^39,40^. Mosquito doesn’t have obligate endosymbionts and relies on commensal bacteria for food digestion, nutrition assimilation, development and reproduction^1,36^. The oversized blood meal female mosquitoes ingested leads to the rapid proliferation of commensal bacteria, which in turn help mosquitoes to digest blood^41^. Tryptophan is one of the essential amino acids that mosquito obtains mainly through blood meals^10^. Majority of tryptophan is metabolized through kynurenine pathway in *Anopheles* mosquitoes as revealed by our LC‐MS analysis. We also show that microbiota participates mosquito Trp metabolism, especially Trp to kynurenine pathway. Accumulation of 3‐HK by depleting microbiota or knockdown of HKT, the 3‐HK catabolizing enzyme, both increase *P. berghei* infection, suggesting that 3‐HK is a key factor that influences the capacity of *An. stephensi* to transmit *P. berghei*. Thus, tryptophan metabolism in mosquitoes is involved in two partners, mosquito and its microbiota. The homeostasis of tryptophan metabolism controlled by these two partners play an important role in determining the vector competence of *An. stephensi*.

3‐HK is toxic by producing reactive radical species under physiological conditions in mosquitoes^11,42^. It is also the substrate of XA that induces the formation of *Plasmodium* microgametocytes^43^. Our results reveal that 3‐HK reduces Per1 expression and compromises PM structure. PM, equivalent to mammal intestinal mucus, is a physical barrier in *Anopheles* mosquitoes that prevents the transmission of *Plasmodium* from gut lumen to epithelium^44,45^. The homeostasis of gut microbiota is essential in PM structure integrity in *Anopheles*, but the underlined mechanism remains unclear^32,33^. Here we provide evidence that the protection of PM by microbiota might through the degradation of PM impairing toxins. Our finding is generally consistent with the mammalian models demonstrating that gut microbiota influences intestinal homeostasis by participating kynurenine pathway^19,42^. A variety of kynurenine metabolites, including Kyn, XA and CA, are the ligands at the aryl hydrocarbon receptor (AHR), a transcription factor that regulates the maturation of different immune cells, epithelial renewal, and barrier integrity, thereby contributing to mucosal homeostasis^21,42,46‐48^. However, the mechanism of the regulation of Per1 expression by 3‐HK remains unclear. It is possible that 3‐HK might influence the expression of *Per1* gene or the stability of Per1 protein. In addition to influencing the expression of major PM protein, 3‐HK might impair PM structure through other unknown mechanisms. XA is a *Plasmodium* exflagellation elicitor and increases *Plasmodium* infectivity in mosquitoes^43,49,50^. However, in our analysis, supplementation of XA at a physiological concentration to mosquitoes fails to promote parasite infection. One possible explanation is that XA fed to mosquitoes might lost its effect when *An. stephensi* is infected with *P. berghei* 24 h later.

The genus *Pseudomonas* is commonly present in multiple mosquito species^4,51^. Several *Pseudomonas* species, including *Pseudomonas aeruginosa, Pseudomonas stutzeri* and *Pseudomonas rhodesiae* reduce pathogen infections in different mosquitoes, but the underlined mechanisms remain not fully understood^51^. Here we demonstrate that *P. alicaligenes* isolated from laboratory reared *An. stephensi* help mosquitoes to defend against *Plasmodium* by promoting 3‐HK catabolism. Kynureninase, KynU, is responsible for 3‐HK degradation^52^. Mutation of this enzyme renders *P. alcaligenes* unable to catabolize 3‐HK, thereby losing the capability to protect PM structure. *An. stephensi* re‐colonized with *KynU* mutant bacterium exhibits reduced inhibitory effect on *P. berghei* infection. However, we failed to detect the downstream product, CA, in vivo possibly due to its low abundance. Nor did we observe any difference of CA levels between microbiota cleared and normal *Anopheles* mosquitoes post blood meal. It is possible that the production of CA is mediated by both mosquitoes and microbiota genes^53^. Elimination of microbiota might facilitate the synthesis of CA by mosquito pathway. Exogenous supplement of either 3‐HAA or CA has no influence on *Plasmodium* infection in *An. stephensi*, further suggesting that *P. alcaligenes* might play a role in converting PM toxic 3‐HK to other inert compounds. Although KynU mutation reduces the inhibitory effect of *P. alcaligenes* on *P. berghei* infection compared to wild type one does, its colonization still increases mosquito resistance to parasites compared to antibiotics treated ones. These results suggest that *P. alcaligenes* plays a dual role in inhibiting *P. berghei* infection, protecting PM by degrading toxic 3‐HK and boosting mosquito immune responses. In summary, our analysis demonstrates how mosquito tryptophan metabolism modulated by gut microbiota influences midgut barrier function, thereby influencing vector competence. Modulating specific tryptophan metabolic pathways in bacteria and mosquitoes might present novel strategies for mosquito control.

## Methods

### Mosquito maintenance and treatments

The *Anopheles stephensi* (Hor strain) was reared under standard laboratory conditions^33^. To eliminate microbiota of *An. stephensi*, newly eclosed mosquitoes were provided with sterile 10% sucrose solution containing penicillin (10 unit/ml), streptomycin (10 µg/ml) and gentamicin (15 µg/ml) for at least four days. To introduce bacteria into the midgut, overnight culture of *P. alcaligenes* were resuspended in 1.5% sterile sucrose solution at a final concentration of 1×10^7^ /ml. A cotton ball soaked with the bacterium was provided to *An. stephensi* for 24 h^54^. *An. stephensi* that starved for 24 h was allowed to feed on *P. berghei* (ANKA) infected BALB/c with parasitemia of 3– 5%^33^. After infection, mosquitoes were maintained at 21°C. Un‐engorged mosquitoes were removed 24 h post blood meal. Midguts were dissected and oocysts were counted microscopically eight days post infection.

### Bacterial strains, genome sequencing and mutation

*Pseudomonas alcaligenes* were isolated and characterized based on the *16S rRNA* sequence from laboratory reared colony. The *16S rRNA* sequences of *Pseudomonas alcaligenes* and reference strains were aligned using the ClustalW multiple alignment tool. The resulting alignments were employed for Neighbor‐Joining phylogenetic reconstructions using MEGA6 software^55^. Genome sequencing of *P. alcaligenes* was performed using Illumina NovaSeq PE150 by the Beijing Novogene Bioinformatics Technology Co., Ltd. After quality control, all good quality paired reads were assembled using the SOAP de novo, SPAdes and ABySS into a number of scaffolds^56‐58^. Coding genes were retrieved by the GeneMarkS program^59^. To predict gene functions, a whole genome blast search (E‐value less than 10^‐5^, minimal alignment length percentage larger than 40%) was performed against KEGG database. The genomic sequence data are available at the National Center for Biotechnology Information’s Sequence Read Archive (accession no. PRJNA686701). The fragment of *P*.*alcaligenes* kynureninase corresponding to bases 61‐1065 was deleted by KnoGen Biotech Ltd, Guangzhou, China. The *P*.*a*.*^ΔKynU^* mutant strain was validated by PCR with the KyuN‐TF/KyuN‐TR primers. The primers were shown in Supplementary Table 5.

### Metabolites Extraction

Extraction of metabolites from mosquitoes was performed according to the previous reference with minor adjustment^60^. Briefly, fifteen mosquitoes (about 30 mg) were pooled for one biological sample. Eight to ten biological replicates were used in the following LC‐MS analysis. Briefly, mosquitoes were snap‐frozen in liquid nitrogen and homogenized in 400 µl of precooled 80% methanol‐ ddH_2_O solution (containing 10 µl of internal standard). After 10 min centrifugation (14,000 g, 4 °C), supernatant was saved and pellet was re‐extracted in 400 µl of precooled methanol‐ddH_2_O two more times. All supernatants were combined and centrifuged at 14,000 g, 4 °C for 10 min, then 1 ml supernatant was transferred into a new tube. Methanol was removed under vacuum using Eppendorf Concentrator plus, and the remaining liquid was lyophilized in a freeze‐drier.

### LC‐MS analysis

Dried metabolite extracts were re‐dissolved in 200 µl of 80% methanol‐ddH_2_O solution for LC‐MS/MS analysis. LC‐MS/MS analysis was performed on a Nexera UHPLC system (Shimadzu, MD) coupled to an ABSciex QTrap 5500 or 6500 mass spectrometer (Framingham, USA). An Agilent ZORBAX RRHD Eclipse Plus C18 (2.1 mm×50 mm, 1.8 µm) column at 35 °C was utilized for LC separation. Samples were injected (1 µl or 10 µl) from the autosampler kept at 4 °C. Then mobile phase A (H_2_O containing 0.1% formic acid) and mobile phase B (acetonitrile containing 0.1% formic acid) were prepared for sample elution. The gradient elution was detailed as follows: 2 % mobile phase B was maintained for 3 min with flow rate setting at 0.3 ml / min and then mobile phase B was increased from 2 % to 80 % within 3 min with flow rate setting at 0.5 ml / min and kept for 2 min, finally the column was reconditioned for 2 min at 2% mobile phase B, the flow rate was set at 0.3 ml / min. For 5500 mass spectrometer detection, mass spectra were acquired on a positive and negative ESI mode with the curtain gas flow of 35 psi, the collision gas of medium, the Ion Spray voltage of 5.5 kV or ‐4.5 kV, the source temperature was 550 °C, the ion source gas 1 (GS1) was 55 psi, the ion source gas 2 (GS2) was 55 psi. Both of the entrance potential (EP) and collision cell exit potential (CXP) was set as 10 V or ‐10 V. For 6500 mass spectrometer detection, mass spectra were acquired on a positive and negative ESI mode with the curtain gas flow of 40 psi, the collision gas of medium, the Ion Spray voltage of 5.5 kV or ‐4.5 kV, the source temperature was 400 °C, the GS1 was 55 psi, the GS2 was 60 psi. Both of the EP and CXP was set as 10 V or ‐10 V. The MRM transition ions of the metabolites and its IS are detailed in Supplementary Table 6. Peak identification and metabolites amount were evaluated based on the known amount of tryptophan metabolites.

### Dietary supplementation of Trp metabolites

Information of tryptophan and tryptophan metabolites used in this study were listed in Supplementary Table 3. Four to six‐day‐old mosquitoes were orally supplemented with 10% sucrose solution containing metabolites with indicated concentration for one day (Supplementary Table 3). Then mosquitoes were starved for 24 h prior to blood feeding.

### RNA extraction and quantitative PCR analysis

RNA was extracted from one mosquito or three pooled midguts using TRIzol reagent (Sigma‐Aldrich, China) according to the standard protocol. Reverse transcription and quantitative PCR were performed as previously described^61^. The *An. stephensi* ribosomal gene *s*7 was used as internal control. Primers were listed in Supplementary Table 5.

### Western Blot analysis

Proteins of 10 midguts 24 h post blood meal were extracted in 300 μl SDS/urea lysis buffer (8 M urea, 2 % SDS, 5 % β‐mercaptoethanol, 125 mM Tris‐HCl). Immunoblotting was performed using standard procedures using the antibodies rabbit anti‐Per1 antibody (1:5000) and mouse anti‐β‐Actin antibody (1:2000) (Abbkine, China). The rabbit polyclonal anti‐Per1 antibody was generated against recombinant Per1 protein (recPer1) corresponding to bases 55‐462 of *per1* CDS (Aste010406) expressed in pET‐42a (Novagen) commercially (GL Biochem Ltd, Shanghai, China).

### Transcriptome analysis

RNA was extracted from 20 pooled midguts dissected from mosquitoes 24 h post infectious blood meal. Three biological replicates of each treatment were used for RNA sequencing. Samples were sent to Majorbio, China for library construction and sequencing using Illumina HiSeq xten. Clean reads were aligned to the reference genome AsteS1.7 (https://www.vectorbase.org/organisms/anopheles-stephensi) using TopHat software^62^. Gene expression was compared using the DESeq2 package in R^63^. Genes with adjusted P‐value less than 0.05 were considered as significantly differentially expressed genes. Raw data are available at the National Center for Biotechnology Information’s Sequence Read Archive (accession no. PRJNA686698).

### Microbiota analysis by 16S rRNA Sequencing

Total DNA was isolated from individual 5‐ day‐ old mosquito using Holmes‐ Bonner method^64^. Six biological replicates were used for the analyzation of population structure of microbiota by 16S rRNA pyrosequencing targeting V3‐ V4 region (341F, 806R) by Majorbio Bio‐Pharm Technology Co. Ltd. (Shanghai, China) using MiSeq platform^65^. A total of 413179 sequences with an average length of 449 bp were obtained. Following quality filtering and chimera sequences removal, sequences analysis were performed by Uparse software^66^. Sequences with ≥ 97% similarity were assigned to the same OTUs and representative sequences were annotated by the Silva Database to identify taxonomic information. Relative abundances were represented by OTU abundances, the number of reads for the given OTU divided by that of total OTUs. The 16S rRNA gene sequences are available at the National Center for Biotechnology Information’s Sequence Read Archive (accession no. PRJNA686689).

### PM structure analysis

Fluorescent immunostaining of Per1 was performed as described^67^. Briefly, the abdomens of *An. stephensi* 45 h post blood meal were collected and fixed in 4% paraformaldehyde at 4 °C overnight. Paraffin embedded sample was sectioned at 5µm thickness and stained with anti‐Per1 (1:100) and Alexa Fluor 546 (1:5000) (Invitrogen). Images were captured using a Nikon ECLIPSE IVi microscope connected to a Nikon DIGITAL SIGHT DS‐U3 digital camera. PM structure was stained by Periodic Acid Schiff (PAS) (Sigma‐Aldrich, China) as describe previously^61^. PM structure was observed under Nikon ECLIPSE IVi microscope connected to a Nikon DIGITAL SIGHT DS‐U3 digital camera.

### TUNEL staining

Fresh prepared slides of the abdomens prior to and 45 h post blood meal were used for TUNEL staining according to the manufacturers’ instructions (Yeasen, Shanghai). Briefly, paraffin sections were dewaxed by xylenes and rehydrated with a graded series of ethanol. Tissue was permeabilized with 20 µg/ml proteinase K for 20 min at room temperature. After PBS rinse, slides were equilibrated with 1× Equilibration Buffer for 30 min at room temperature, then equilibration Buffer was removed and slides were incubated with TUNEL reaction mixture that contains Alexa Fluor 488‐12‐dUTP at 37 °C for 1 hr. Apoptosis positive signal was acquired with 488 nm excitation using a Nikon ECLIPSE IVi microscope. Nuclei were stained with DAPI.

### DHE staining

The mosquito midguts were dissected in PBS at 0 h and 24 h post normal blood meal. DHE staining was performed as previously described^68^. Briefly, midguts were incubated with 5 μM dihydroethidium (DHE) (Beyotime, China) at room temperature for 20 min in the dark. Image was captured using Zeiss‐LSM880 confocal microscope.

### PH3 staining

PH3 staining was performed as previous described^69^. Briefly, midguts were dissected in PBS at 0 h and 24 h post normal blood meal. Dissected guts were fixed with 4% paraformaldehyde for 20 min at room temperature, then rinsed in PBT with 0.1% Triton X‐100. After blocking with 3% BSA for 2 h, midguts were incubated with 1:1000 anti‐PH3 (Merck, Germany) overnight at 4°C. Alexa Fluor 546 (1:5000) (Invitrogen) was used as secondary antibody. Images were acquired by Zeiss‐LSM880 confocal microscope.

### Statistics

All statistical analyses were performed using GraphPad Prism software. The details of statistical methods are provided in the figure legends. Difference of oocyst number between two groups was analyzed using the Mann‐Whitney test, among more than two groups was analyzed using ANOVA. Difference of metabolites between two groups was analyzed using the Student’s t‐test, among more than two groups was analyzed using ANOVA. The statistical analysis of metabolomics data and transcriptome data are described in the corresponding methods.

## Acknowledgements

We thank Guoliang Zhang, Chuyang Wang and Professor Jinying Gou at School of Life Sciences, Fudan University for their technical advice on metabolites analysis. This work was supported by the National Natural Science Foundation of China (U1902211, 31822051) and the National Institutes of Health Grant (R01AI129819) to J. W.

## Author contributions

Y.F., Y. P., J.W. and H.T. designed experiments, interpreted results and wrote the paper. Y. P. and Y. A. performed metabolites analysis. H. W. and X. S. conducted PM structure analysis experiments. H. W. and Y. F. conducted bacteria recolonization and *Plasmodium* infection experiments. Y.F. conducted and analyzed results from all additional experiments. J. W. and H.T. supervised the study. All authors discussed the results and commented on the manuscript.

---

**Extended Data Fig. 1.**
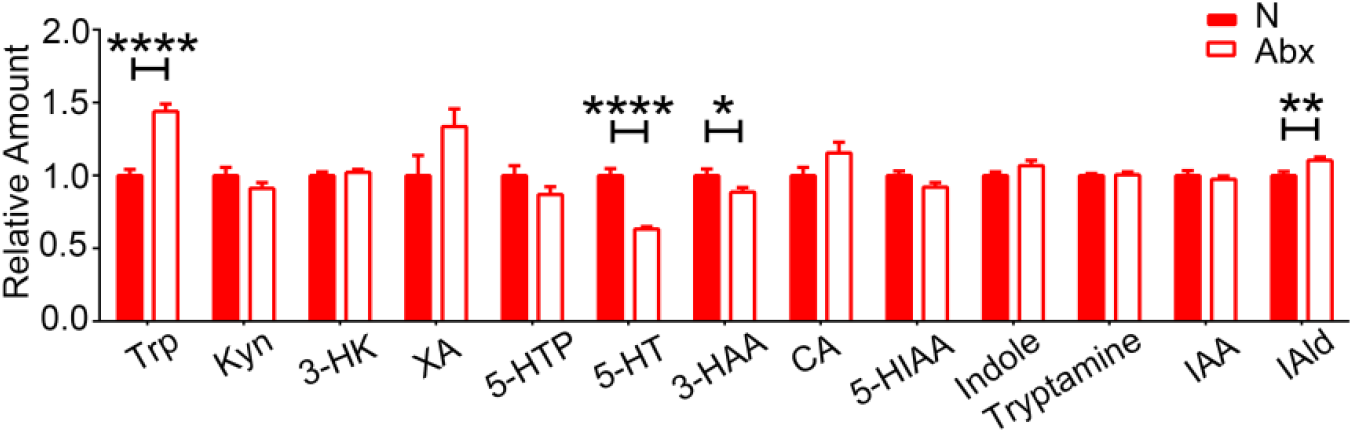
The relative amounts of Trp metabolites in normal (N) and antibiotics‐treated (Abx) mosquitoes 24 h post normal blood meal. Error bars indicate standard errors (n=10). Significance was determined by Student’s *t*‐test. * P<0.05, **P<0.01, ****P<0.0001.

**Extended Data Fig. 2.**
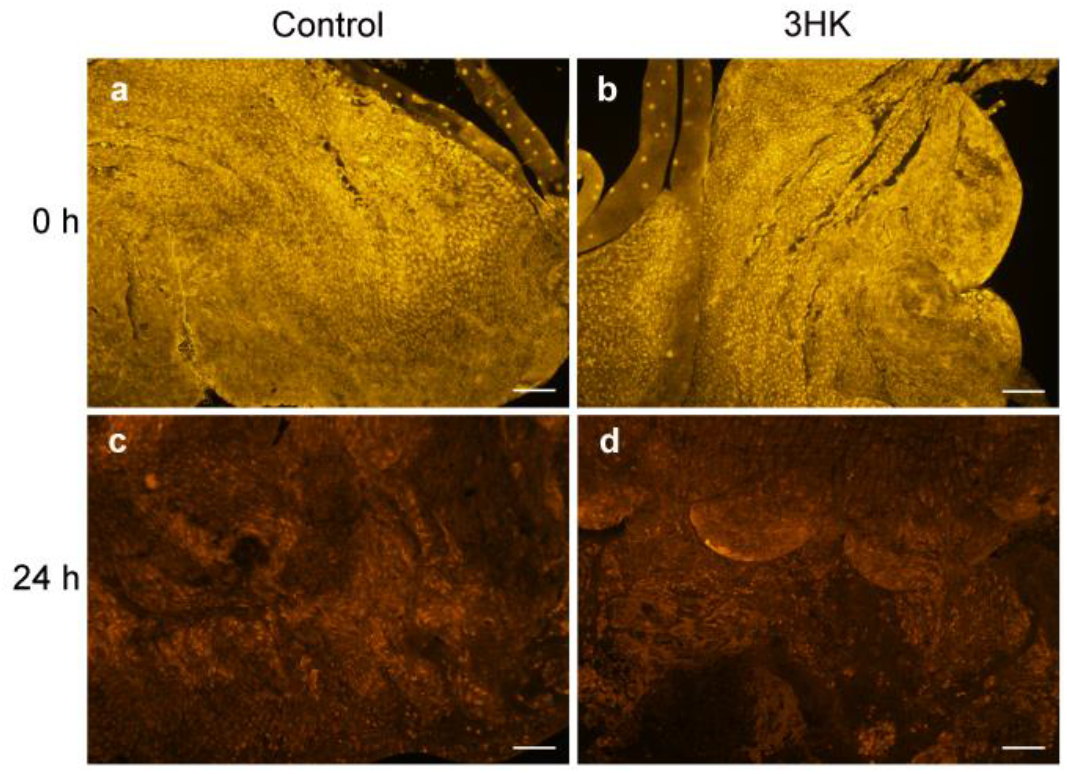
DHE staining of midguts in 3‐HK‐treated (‐) and control (+) mosquitoes at 0 h (**a, b**) and 24 h (**c, d**) post normal blood meal. Scale bars represent 100 µm. Images are representative of at least eight midguts.

**Extended Data Fig. 3.**
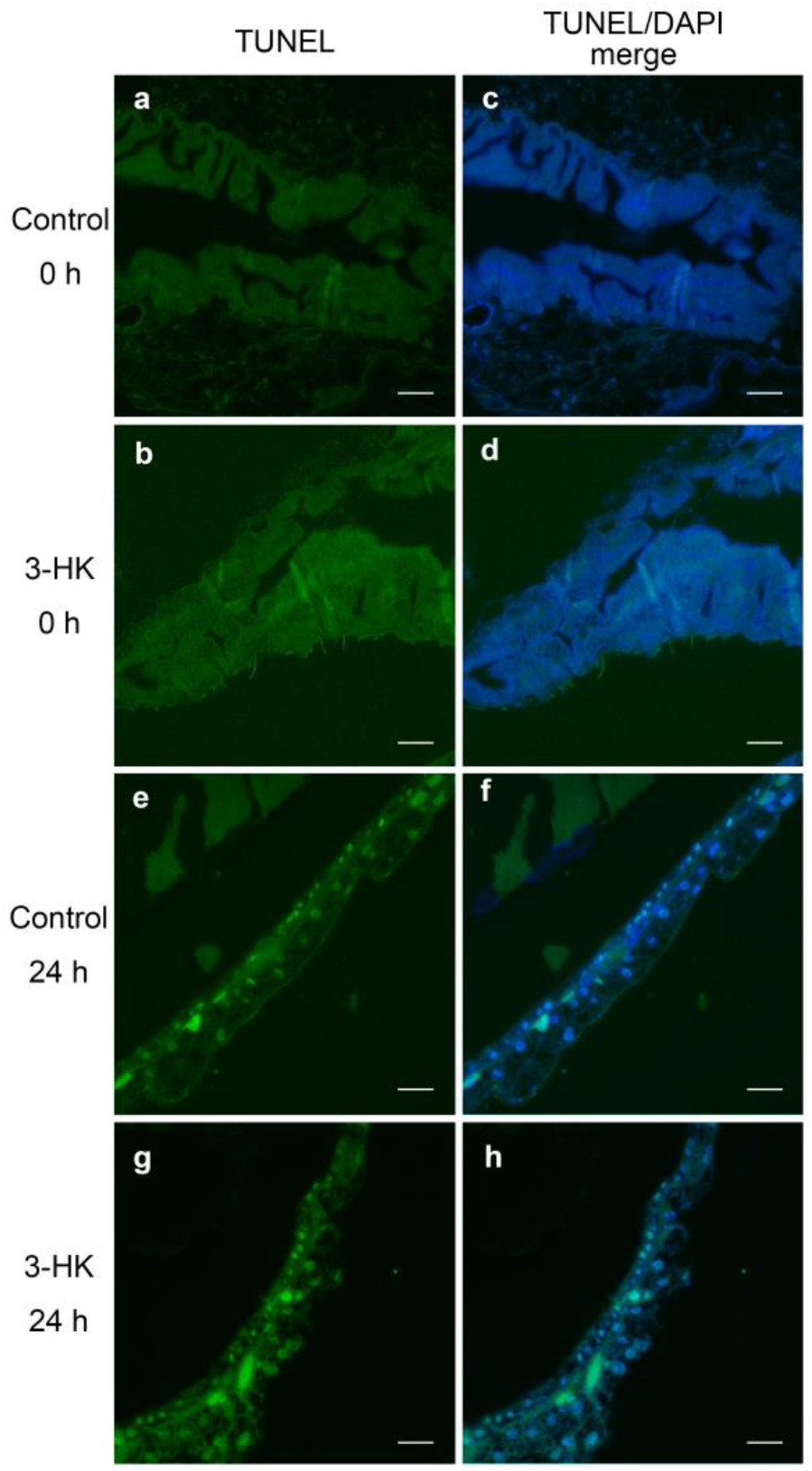
TUNEL staining of midgut from 3‐HK‐treated mosquitoes and controls at 0 h (**a‐d**) and 24 h (**e‐i**) post normal blood meal. Scale bars represent 50 µm. Images are representative of at least four midguts.

**Extended Data Fig. 4.**
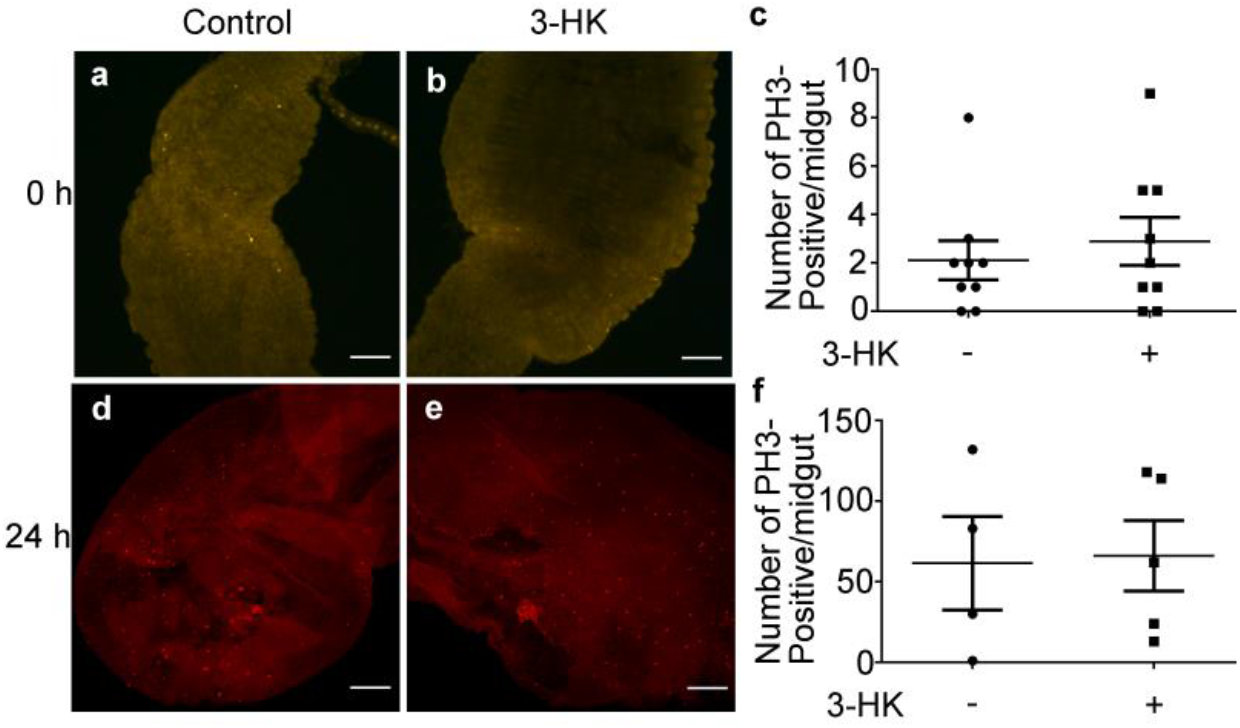
PH3 staining of midguts from 3‐HK‐treated mosquitoes and controls at 0 hr (**a, b**) and 24hr (**d, e**) post normal blood meal. Scale bars represent 100 µm. Quantification of PH3‐positive cells are shown in (**c**) and (**f**). Error bars indicate standard errors. Significance was determined by Student’s *t*‐test.

**Extended Data Fig. 5.**
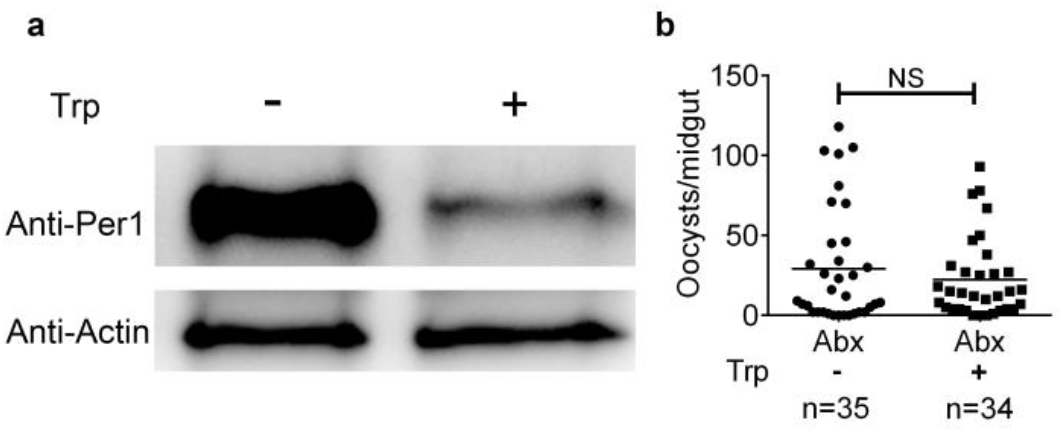
Trp supplementation impairs PM structure. **a,** Western blot of Per1 in the midgut of Trp‐treated (+) mosquitoes and control (‐) 24 h post normal blood meal. **b**, Effect of Trp on *P. berghei* infection in Abx mosquitoes. Data were pooled from two independent experiments. Each dot represents an individual mosquito. Horizontal black bars indicate the median values. Significance was determined by Mann‐Whitney tests.

**Extended Data Fig. 6.**
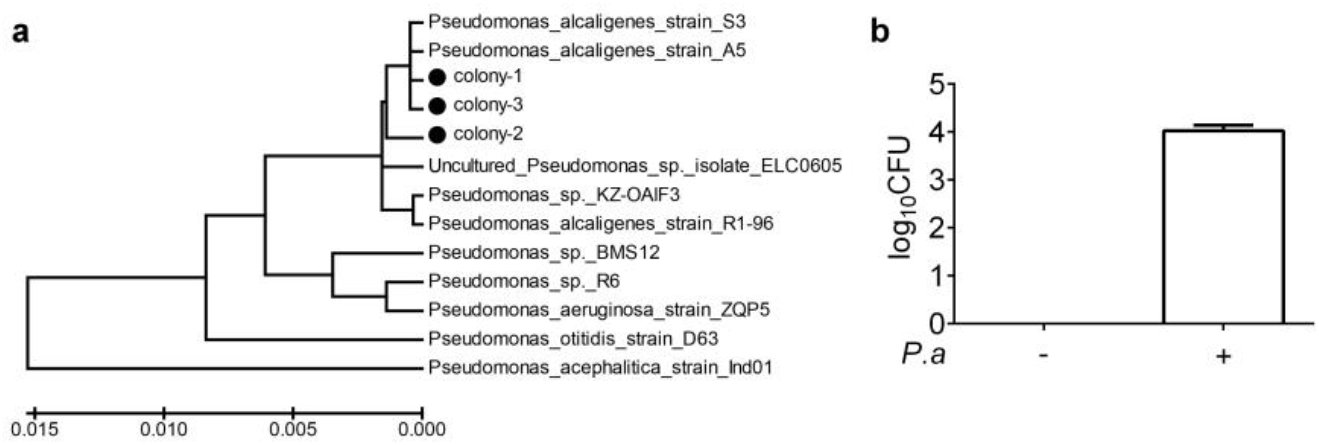
Characterization of culturable *Pseudomonas* sp. in *An. stephensi*. **a,** Phylogenetic tree showing the relationship between *P. alcaligenes* and other *Pseudomonas* sp. based on *16S rRNA* genes sequences. **b**, Bacterial loads in the midgut of Abx (‐) and *P. alcaligenes* (+) recolonized mosquitoes before blood meal (n=6). Error bars indicate standard errors.

**Extended Data Fig. 7.**
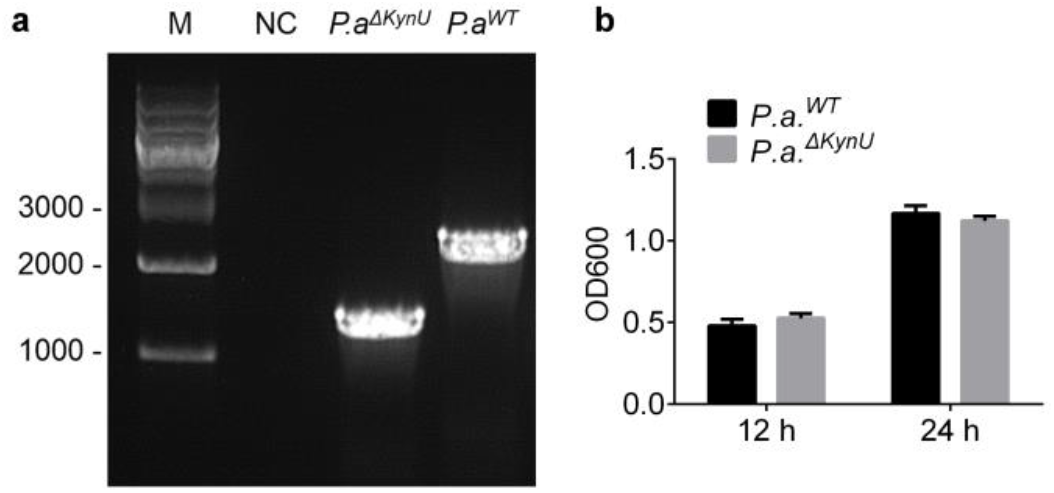
Mutation of KynU in *P. alcaligenes*. **a,** Confirmation of *KynU* deletion by PCR. **b**, The growth rate of *P*.*a*.^*ΔKynU*^ and *P*.*a*.^*ΔKynU*^ *in vitro*.

